# A probabilistic approach to discovering dynamic full-brain functional connectivity patterns

**DOI:** 10.1101/106690

**Authors:** Jeremy R. Manning, Xia Zhu, Theodore L. Willke, Rajesh Ranganath, Kimberly Stachenfeld, Uri Hasson, David M. Blei, Kenneth A. Norman

## Abstract

Recent research shows that the covariance structure of functional magnetic resonance imaging (fMRI) data - commonly described as *functional connectivity* - can change as a function of the participant’s cognitive state (for review see [35]). Here we present a Bayesian hierarchical matrix factorization model, termed *hierarchical topographic factor analysis* (HTFA), for efficiently discovering full-brain networks in large multi-subject neuroimaging datasets. HTFA approximates each subject’s network by first re-representing each brain image in terms of the activities of a set of localized nodes, and then computing the covariance of the activity time series of these nodes. The number of nodes, along with their locations, sizes, and activities (over time) are learned from the data. Because the number of nodes is typically substantially smaller than the number of fMRI voxels, HTFA can be orders of magnitude more efficient than traditional voxel-based functional connectivity approaches. In one case study, we show that HTFA recovers the known connectivity patterns underlying a collection of synthetic datasets. In a second case study, we illustrate how HTFA may be used to discover dynamic full-brain activity and connectivity patterns in real fMRI data, collected as participants listened to a story. In a third case study, we carried out a similar series of analyses on fMRI data collected as participants viewed an episode of a television show. In these latter case studies, we found that the HTFA-derived activity and connectivity patterns can be used to reliably decode which moments in the story or show the participants were experiencing. Further, we found that these two classes of patterns contained partially non-overlapping information, such that decoders trained on combinations of activity-based and dynamic connectivity-based features performed better than decoders trained on activity or connectivity patterns alone. We replicated this latter result with two additional (previously developed) methods for efficiently characterizing full-brain activity and connectivity patterns.

## Introduction

The most common approaches for analyzing functional Magnetic Resonance Imaging (fMRI) data involve relating, in individual images, the activity of individual voxels or multi-voxel spatial patterns of brain activity to the subject’s cognitive state [14, 15, 27, 41]. In contrast, functional connectivity analyses correlate the time series of activities *across images* of pairs of voxels [30]. Functional connectivity analyses have already led to new insights into how the brain’s correlational structure changes during different experimental conditions [35].

The size of the full-brain functional connectivity matrix grows with the square of the number of voxels. Because of this rate of growth, filling in its entries and storing it in memory can become intractable for fMRI images with tens of thousands of voxels. For example, for a series of 50,000 voxel images, each connectivity matrix occupies approximately 5 GB (storing only the upper triangle, using single precision floating point entries). Storing many such matrices in memory (e.g. to compare different subjects and/or experimental conditions) can exceed the limits of modern hardware. Further, many of the algorithms used to relate multivariate patterns of voxel activities in individual images to cognitive states or experimental conditions (e.g. [22, 27]) use superlinear time and memory with respect to the number of features; this computational expense makes it impractical to use the same techniques to examine correlational data (although see [37] for another promising approach using massive parallelization on a specialized computing cluster).

Previous work circumvents this issue by using functionally [12, 20, 29, 39] or anatomically [3, 13, 31] defined voxel clusters or regions of interest. These approaches segment the brain into discrete components, and then examine interactions or correlations between the activity patterns exhibited by those components (rather than attempting to examine every voxel-to-voxel interaction). In other words, these approaches encapsulate an intuition about how our brains work-specifically, that our brains are composed of a small number of network nodes that interact with each other.

One such class of methods entails pre-selecting a small number of regions of interest (ROIs; e.g. motor cortex; [5]) or a seed voxel (e.g. a single voxel within the posterior cingulate; [21]). This procedure reduces the connectivity matrix from a *V* × *V* matrix (where *V* is the total number of voxels in each brain volume) to a much smaller *V*_ROI_ × *V* matrix (where *V*_ROI_ is the number of seed or ROI voxels). However, reducing the connectivity matrix in this way precludes finding connectivity patterns unrelated to the ROI or seed region. For example, if the analysis is limited to connectivity patterns between motor cortex and the rest of the brain, this precludes finding patterns of connectivity that do not involve the motor cortex (e.g. connectivity between prefrontal cortex and the hippocampus). Other work has examined full-brain connectivity patterns via ROI-to-ROI connectivity matrices [e.g. as defined using anatomical data; 24]. This method provides a tractable means of examining full-brain connectivity patterns by assuming that each ROI is perfectly uniform and discrete.

A related technique that does not require pre-selecting ROIs or seed regions is to compute the full voxel-to-voxel connectivity matrix, and then to threshold the connection strengths such that one only examines the most reliable connections [7]. This approach yields a sparse voxel-to-voxel connectivity matrix that may be efficiently manipulated (provided that it is sufficiently sparse). One drawback to this approach is that it is not always clear how to set the connectivity strength threshold; for example, setting too high a threshold will leave out potentially important patterns, whereas setting too low a threshold will not substantially reduce the computational burden (as compared with examining the original voxel-to-voxel connectivity matrix). However, regularization techniques such as sparse regression [e.g., 36] show promise. Another potential drawback is that it is not clear that the strongest connections are necessarily the most informative; for example, a sub-threshold connection may still carry cognitively relevant information.

Other approaches have focused on reducing the dimensionality of the connectivity patterns (see [38] for a comparison between threshold-based and dimensionality-reduction-based approaches). For example, clustering-based approaches attempt to group together voxels into “factors” that exhibit similar activity patterns over time, across subjects [11]. One may then examine connectivity patterns between the factors rather than between the voxels [this approach was originally developed in the positron emission tomography literature using Principle Components Analysis (PCA) [16]].

Here we propose *hierarchical topographic factor analysis* (HTFA), a Bayesian factor analysis model specifically designed to facilitate analyses of brain network dynamics. HTFA is focused on finding spherical nodes that can be used to efficiently explore, analyze, and understand full-brain network dynamics. Like other dimensionality reduction methods, HTFA provides a compact means of representing full brain connectivity patterns that scales well to large datasets [1]. But HTFA goes beyond these methods: (a) it provides a natural means of determining how many network nodes should be used to describe a given dataset; (b) it allows those nodes to be overlapping rather than forcing nodes to be fully distinct; and (c) it constrains the nodes to be in similar (but not necessarily identical) locations across people. Further, HTFA decomposes brain images into sums of spatial functions, which supports seamless mapping between images of different resolutions (e.g. different voxel sizes) and potentially between different recording modalities (e.g. fMRI, EEG, ECoG, MEG). We return to these latter points in the *Discussion*.

HTFA casts each subjects’ brain images as linear combinations of latent factors [Gaussian radial basis functions (RBFs)]. Each RBF can be interpreted as a spherical *node* in a simplified representation of the brain’s networks. (The number of nodes, *K*, is determined from the data.) In this way, HTFA is a spatial model: these nodes reflect structures localized in 3D space whose activity patterns influence the observed voxel activities. This is conceptually different from approaches that interpret voxel activities directly: HTFA defines an explicit model for how voxels relate to each other according to their relative locations in space (see also [18]). One advantage of constraining HTFA’s nodes to be RBFs is that they are spatially compact. In this way, HTFA is qualitatively similar to spatial ICA [6]. However, unlike spatial ICA, HTFA further constrains nodes to be in similar locations across subjects, providing a natural means of combining or comparing connectivity across individuals. (ICA with dual regression [2] also models hierarchical patterns across participants.) Note that in this paper we have constrained the RBF nodes to be isotropic, but in principle this assumption could be relaxed to support any parameterizable spatial function (e.g., non-isotropic Gaussians, mixtures of Gaussians, 3D wavelets, etc.). These spatial functions could also be constructed to incorporate detailed conductance models (e.g. informed by diffusion tensor imaging data). We have left these extensions to be explored in future work.

Applying HTFA to an fMRI dataset reveals the locations and sizes of these network nodes (i.e. the centers and widths of their RBFs), as well as the per-image node weights. If a given subject has contributed *N* images to the dataset, then the subject’s *N* by *K* node weights matrix may be viewed as a low-dimensional embedding of their original data. Further, the pairwise correlations between columns of this weight matrix reflect the signs and strengths of the node-to-node connections (just as the pairwise correlations between voxel time series reflect the corresponding “connectivity” in voxel-based functional connectivity analyses).

The next section provides a descriptive overview of the HTFA model. *Materials and methods* describes how we efficiently fit the model to a multi-subject fMRI dataset using *maximum a posteriori* inference. The *supplemental materials* contain a complete formal (mathematical) description of the model. To validate our approach, we first generated a set of 50 synthetic datasets for which the underlying activity patterns, node locations and sizes, etc. were known, and we show that HTFA recovers these known patterns. Then we applied HTFA to two (real) fMRI datasets collected as participants listened to an audio recording of a story (Case Study 2) and watched an episode of a television show (Case Study 3). We show that the moment-by-moment patterns uncovered by HTFA may be used to decode which moments in the story or show the participants were experiencing.

## Model description

### Overview

HTFA is a member of a family of models, called *factor analysis* models, that includes Topographic Factor Analysis (TFA) [26], Topographic Latent Source Analysis (TLSA) [18], Principal Components Analysis (PCA) [28], Exploratory Factor Analysis (EFA) [33], and Independent Components Analysis (ICA) [10, 25], among others. If we have organized our collection of images (from a single subject) into an N by V data matrix Y (where N is the number of images and V is the number of voxels), then factor analysis models decompose Y as follows:

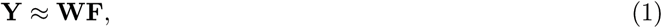

where **W** is an *N* by *K* weight matrix (which describes how each of *K* factors are activated in each image), and **F** is a *K* by *V* matrix of factor images (which describes how each factor maps onto the brain). Note that, in the general case, this decomposition is underspecified-in other words, there are infinitely many solutions for **W** and **F** that approximate the data equally well. What differentiates factor analysis models is the particular constraints they place on what form **W** and/or **F** should take (i.e. by changing the function being optimized in order to settle on a specific choice of **W** and **F**). We may then use **W** as a low-dimensional embedding of the original data (e.g. to facilitate interpretability or improve computational tractability), or we may choose to examine the factor images in **F** to gain insights into the spatial structure of the data.

In related approaches such as PCA and ICA, the entries of **W** and **F** can, in principle, be any pattern of real numbers. In PCA, each row of **F** is an eigenvector of the data covariance matrix, and **W** is chosen to minimize the reconstruction error (i.e. to make **WF** as close as possible to **Y** in terms of mean squared error). In ICA, the goal is to minimize the statistical dependence between the rows of **F** while also adjusting **W** to minimize the reconstruction error. In this way, the factor images (the rows of F) obtained using PCA and ICA are unstructured images (i.e. activity patterns) of the same complexity as individual observations in the original dataset: each factor is parameterized by *V* numbers (1 parameter per voxel).

In TFA (and TLSA), each row of **F** is parameterized by the center parameter, *μ*, and the width parameter, *λ*, of an RBF. If an RBF has center *μ* and width *λ*, then its activity RBF(**r**|*μ*, *λ*) at location **r** is:

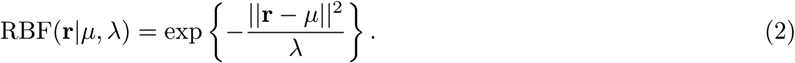

The factor images are filled in by evaluating each RBF, defined by the corresponding parameters for each factor, at the location of each voxel. In contrast to the factors obtained using PCA or ICA, TFA’s more constrained factors may be represented much more compactly; each factor corresponds to the structure or group of structures in the brain over which the factor spreads its mass (which is governed by *μ* and *λ*). As highlighted above, TFA’s factors may be conceptualized as nodes located in 3D space whose activity patterns influence the observed brain data.

HTFA works similarly to TFA, but places an additional constraint over the nodes to bias all of the subjects to exhibit similar nodes. Whereas TFA attempts to find the nodes that best explain an individual subject’s data, HTFA also attempts to find the nodes that are common across a group of subjects (Fig. 1). This is important, because it allows the model to jointly consider data from multiple subjects.

**Figure 1.**
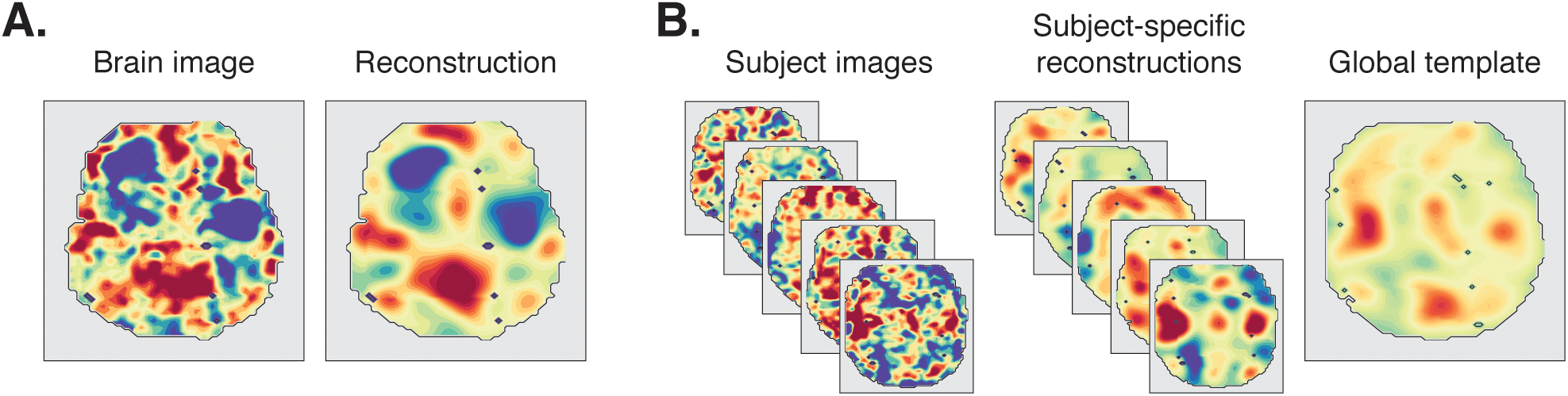
Hierarchical topographic factor analysis. **A. A brain image and its associated reconstruction.** The left sub-panel displays a single horizontal slice from a single subject; the right sub-panel displays its associated HTFA reconstruction, which we obtain by summing together the weighted images of the subject’s RBF nodes. **B. Explaining data across subjects.** The left and middle sub-panels display example images from five subjects (left sub-panel), and their associated reconstructions (middle sub-panel). The right sub-panel displays the approximation of all of the single-subject images in the left sub-panel, obtained by setting the weights of the global template’s nodes to their average weights in the subject-specific reconstructions. The locations of the RBFs in the global template reflect commonalities across subjects, whereas the single-subject RBF locations reflect the associated subject-specific idiosyncrasies.

HTFA handles multi-subject data by defining a *global template*, which describes in general where each RBF is placed, how wide it is, and how active its node tends to be. In addition to estimating how nodes look and behave in general (across subjects), HTFA also estimates each individual’s *subject-specific template*, which describes each subject’s particular instantiations of each RBF (i.e. that subject’s RBF locations and widths) and the node activities (i.e. the activities of each of that subject’s RBF nodes in each of that subject’s images). These node activities, in turn, may be used to estimate each subject’s full-brain functional connectivity patterns. Further, because the subject-specific templates are related to each other (hierarchically [17], via the global template), a given node’s RBF will tend to be located in about the same location, and be about as large, across all of the subject-specific templates. Because each subject has the same set of nodes (albeit in slightly different locations and with slightly different sizes) we can run analyses that relate node activity across subjects (as in the inter-subject functional connectivity analyses described below).

## Materials and methods

### Applying HTFA to multi-subject fMRI datasets

We use a *maximum a posteriori* (MAP) inference procedure to compute the most probable RBF nodes and node weights. The procedure has three basic steps: initialization (during which we set the prior over each node’s parameters); fitting subject-specific node parameters for each subject (given the prior); and updating the global template (using the subject-specific parameters). When we carry out the full inference procedure, we first perform a cross-validation step (described in the next sub-section) to determine the optimal number of nodes, *K*. We then randomly initialize *K* RBF nodes by drawing their parameters from the prior distribution (see *supplemental materials*). Finally, we iterate between updating the subject-specific parameters (using the current global template as the prior) and the global template (using the latest estimates of the subject-specific parameters) until the largest change in any parameters value from the previous iteration to the current iteration is less than a pre-determined threshold value, *ϵ*. (We typically set *ϵ* to be the length of the longest voxel dimension.) Once the global centers and widths have converged, we run an additional step whereby we re-compute the per-image node weights for each subject. We developed an efficient implementation of our algorithm for applying HTFA to large multi-subject fMRI datasets [1]; additional details may be found there. We also published a Python toolbox that enables users to apply HTFA to multisubject fMRI datasets [8].

#### Estimating the optimal number of nodes

When *K* (the number of nodes) equals *V* (the number of voxels), HTFA can exactly recover the data by setting the RBF widths to be very small and the per-image node activities equal to the voxel activities. Therefore setting *K* = *V* represents one logical extreme whereby HTFA loses no information about the original data but also achieves no gains in how efficiently the data are represented. At the other extreme, when *K* = 1, HTFA achieves excellent compression but poor reconstruction accuracy (only a single activity value may be represented for each image). In practice, we will typically want to set *K* to some value between these extremes. Specifically, we want to choose the minimum *K* that is expected to explain the data up to a pre-defined level of precision, *q*.

Given a multi-subject dataset, we first select *s*_*training*_ subjects at random (from the full set) to participate in the cross validation procedure. For each of those subjects, we select (at random) a set of *n*_*training*_ images from each subject’s data (here we set this number to be 70% of the subject’s images). We fit HTFA to these randomly selected images to estimate the subject-specific node centers (*μ*_1…*K*,1…*S*_) and widths (*λ*_1…*K*,1…*S*_). Next, of the remaining *n*_*test*_ = *N*_s_ - *n*_*training*_ images for each subject, we select (at random) 70% of the voxels to estimate the per-image node weights, *w*_1…*N*,1…*K*,1…*S*_. Finally, we use the estimated centers, widths, and weights to reconstruct the voxel activities for the remaining 30% of the voxels in those *n*_*test*_ images. The mean squared error between the reconstructed and true (observed) voxel activities provides an error signal that we can use to optimize *K*. In particular, starting from a minimum value of *K* = *δ*_*K*_, we use the above procedure to compute the mean squared error for *δ*_*K*_. We then increase *K* (in increments of *δ*_*K*_) until the mean squared error is less than our pre-defined threshold, *q*. (In this paper we set *δ*_*K*_ = 100 and *q* = 0.25.)

### Inferring dynamic full-brain inter-subject functional connectivity patterns

Inference yields, for each subject, an *N*_*s*_ by *K* matrix, **W**_*s*_, of per-image node weights (i.e. activity patterns). We can estimate the functional connectivity between each pair of nodes by computing the correlation between the columns of **W**_*s*_. This approach is analogous to standard voxel-wise techniques for estimating functional connectivity [5]. Further, because the columns of **W**_1…*S*_ correspond to the same nodes across the different subjects, since all of the nodes are linked through the global template, the set of these weight matrices provide a convenient means of testing hypotheses related to the connectivity strengths.

In our analyses for Case Studies 2 and 3, we used *inter-subject functional connectivity* [ISFC; 32] to isolate the time-varying correlational structure (functional connectivity patterns) that was specifically driven by the story participants listened to (Case Study 2) and television show participants watched (Case Study 3). We first applied HTFA to the fMRI datasets to obtain a time series of node activities for every participant (where *K* = 700 for Case Study 2 and *K* = 900 for Case Study 3; see *Estimating the optimal number of nodes*). We obtained ISFC matrices for each of a series of overlapping temporal windows (*sliding windows*). To do so, we performed the following analysis for each sliding window (which contained a 90 s time series of node activities for each node and participant). For each participant, we computed the correlation between the activities of each node from that participant (during that sliding window) and the average activities of every node during the same window (where the average was taken across all of the other participants). The result, *C*_*s,t*_ was a *K* by *K* correlation matrix for a single participant (*s*), during a single sliding window (*t*). We computed the ISFC matrix (across participants) during time *t* as:

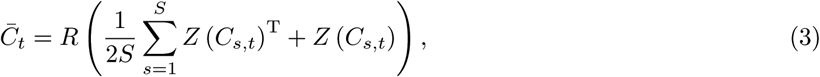

where *Z* is the Fisher *z*-prime transformation [40]:

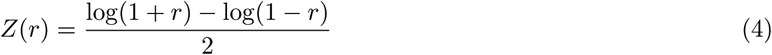

and *R* is the inverse of *Z*:

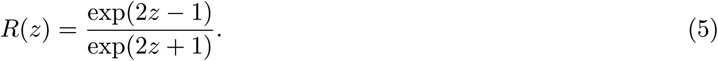

For additional details and discussion of ISFC see [32].

## Decoding analysis

We asked whether the moment-by-moment HTFA-derived patterns we identified in Case Studies 2 and 3 were reliably preserved across participants, and (in Case Study 2) whether the degree of agreement across participants was modulated according to the cognitive salience of the stimuli participants experienced. For example, prior work has shown that different participants exhibit similar responses while experiencing richly structured stimuli (such as story listening), whereas participants exhibit less stereotyped responses while experiencing less structured stimuli (such as resting with their eyes open in the scanner) [32].

To study these phenomena, we randomly divided the Case Study 2 participants into two groups, for each experimental condition: intact (i.e. participants who listened to the original story recording), paragraph-scrambled (i.e. participants who listened to an altered recording where the paragraphs occurred in a randomized order), word-scrambled (i.e. participants who listened to an altered recording where the words occurred in a randomized order), and rest (i.e. participants who rested with their eyes open in the scanner, without listening to any story). Participants within each condition experienced the same auditory stimuli, but the cognitive salience (i.e. how meaningful the stimuli were) varied systematically across these experimental conditions. (For the experiment presented in Case Study 3, every participant experienced the analog of the “intact” condition.)

For each experimental condition, we computed the mean voxel, node, or factor activities within each sliding window (this resulted in either a *V*-dimensional or *K*-dimensional vector for each moment of the story). For each group of participants in turn, we compared these activity patterns (using Pearson correlations) to estimate the story times each pattern corresponded to. In the activity-based analyses shown in Figure 4 (darker bars in each color group), we used these activity vectors to decode which moments of the story participants were listening to. Specifically, we asked, for each sliding window (*t*): what are the correlations between the first group’s activity pattern at time *t* and the second group’s activity patterns in *every* sliding window (this yielded one correlation value per sliding window). We used the best-matching pattern (i.e. the activity pattern with the strongest positive correlation) to estimate which story time sliding window *t* corresponded to.

**Figure 4.**
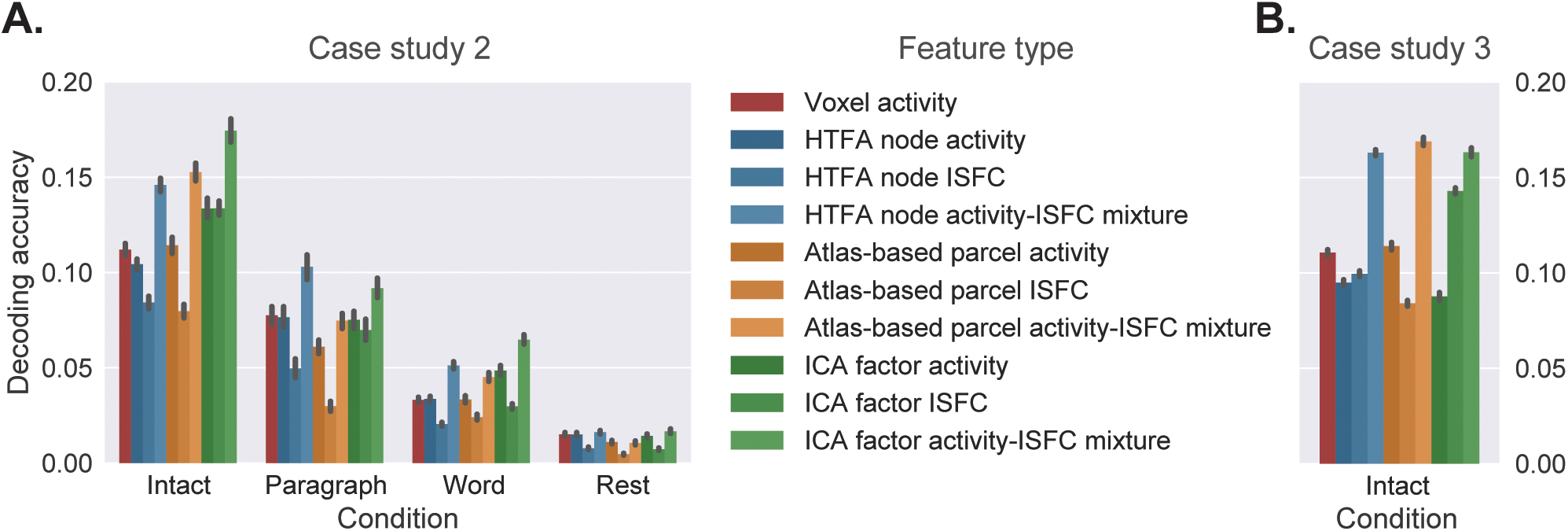
Decoding accuracy. Bars of each color display cross-validated decoding performance for decoders trained using different sets of neural features: whole-brain patterns of voxel activities (red); HTFA-derived node activities (blue); anatomically derived parcel activities [31] (yellow); and dual regression ICA-derived factor activities (green). The shading reflects post-processing applied to each class of feature prior decoding: activity-based decoding (dark shading); ISFC-based decoding (medium shading); and a 50-50 mixture of activity-based and ISFC-based decoding (light shading). Panel **A.** displays the decoding performance for Case Study 2 and panel **B.** displays the decoding performance for Case Study 3. All error bars denote bootstrap-estimated 95% confidence intervals, estimated using 5000 iterations.

We used a similar approach to examine moment-by-moment ISFC patterns for each of the two groups of participants (i.e. for each condition, we obtained one ISFC pattern for each sliding window, for each of the two groups). For the ISFC analysis shown in Figure 4 (medium-shaded bars in each color group), we reshaped these ISFC patterns into vectors, and used the same correlation-based technique to label each group’s sliding windows according to how well they matched the ISFC patterns in the other group’s sliding windows.

Finally, we carried out a mixed activity-based and ISFC-based decoding analysis by combining the estimates of the two above decoders. Specifically, for each sliding window (from one group of participants), we computed the correlations between the activity patterns from each sliding window from the other group, and the correlations between the ISFC patterns from each sliding window of the other group. We used the average of these two correlations to label each group’s timepoints in the “mixed” decoding analysis shown in Figure 4 (lightly shaded bars in each color group). In Figure 5 we extended this analysis by changing the relative weights of the activity-based and ISFC-based decoders using a mixing parameter, *ϕ*, where *ϕ* = 0 corresponds to activity-based decoding; *ϕ* = 1 corresponds to ISFC-based decoding; and 0 < *ϕ* < 1 reflects a weighted mixture of activity-and ISFC-based decoding. (Using this notation, we fixed *ϕ* = 0.5 for all mixture analyses in Fig. 4.)

**Figure 5.**
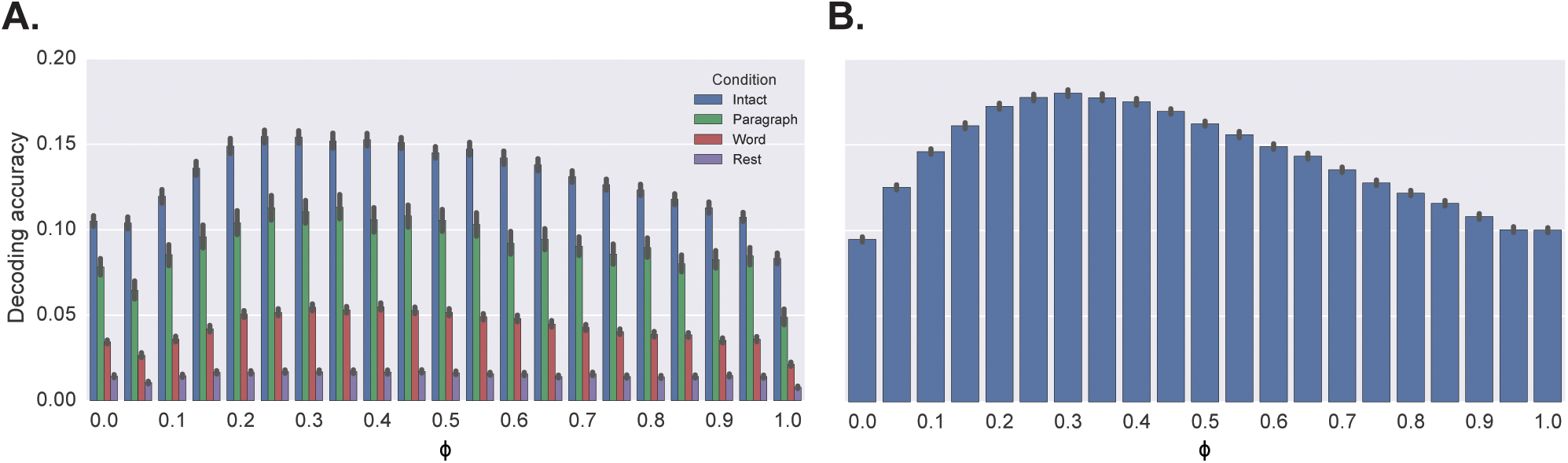
Decoding accuracy as a function of mixing proportion (*ϕ*). The bar heights indicate the decoding accuracy achieved for each value of the mixing parameter, *ϕ*, where *ϕ* = 0 corresponds to HTFA node activity-based decoding; *ϕ* = 1 corresponds to ISFC-based decoding; and 0 < *ϕ* < 1 reflects a weighted mixture of activity-and ISFC-based decoding. Panel **A.** displays the decoding performance for Case Study 2 (colors denote the experimental condition) and panel **B.** displays the decoding performance for Case Study 3.

We note that the decoding test we used is more conservative than those used in some previously reported timepoint decoding studies (e.g. [23]) because we count a timepoint label as incorrect if it is not an exact match, even if it overlaps substantially with the correct label. For example, if our decoder matches the 0-60 TR window from group 1 with the 1-61 TR window from group 2 (i.e. 88.5 of the 90 seconds are overlapping), our performance metric considers this to be a decoding failure, indistinguishable (performance-wise) from if the group 1 and group 2 windows had not overlapped at all; by contrast, [23] used a more liberal procedure where they only compared the correct time window (e.g. 0-60 TR) to the exactly matching window or non-overlapping time windows (e.g. 61 - 120 TR), but not to partially overlapping windows (e.g. 1-61 TR). We chose to use this conservative test because our decoders attained over 99% accuracy for both voxel-based and HTFA-derived neural features (on data from the intact condition of the experiment in Case Study 2, as well as in Case Study 3) when we used the more liberal “standard” matching procedure. This made it difficult to achieve our goal of comparing and distinguishing between decoders trained on different neural features, hence our more conservative test.

## Results

We applied HTFA to three types of data. In Case Study 1, we applied HTFA to synthetic data, and in Case Studies 2 and 3 we applied HTFA to fMRI data from human participants.

### Case study 1: examining network recovery from synthetic data

We generated 50 synthetic datasets, each comprising 200 brain images from each of 10 simulated subjects. Each image volume was a 50 x 25 x 25 rectangular block of 31,250 1x2x2 mm voxels. To generate the voxel activities in each image, we randomly placed 20 RBF centers in a “template” brain volume (drawn randomly without replacement from the set of 31,250 voxel locations), and randomly assigned a positive width to each RBF (where log(width) 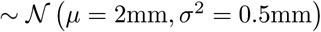). We then generated each subject’s RBF centers and widths by adding a small amount of Gaussian noise to the corresponding parameters in the template (for each parameter dimension, noise 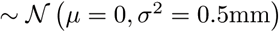). Next, we set the time-varying activities of each simulated subject’s RBF nodes to exhibit a pre-defined sequence of activity patterns. Specifically, the node activations were defined by a symmetric Toeplitz matrix whose first row was the sequence (1, 2,…, 19, 20, 20,19,…, 2,1), tiled 5 times to yield an activation for each node in each of the 200 images (Fig. 3D, left panel).

**Figure 3.**
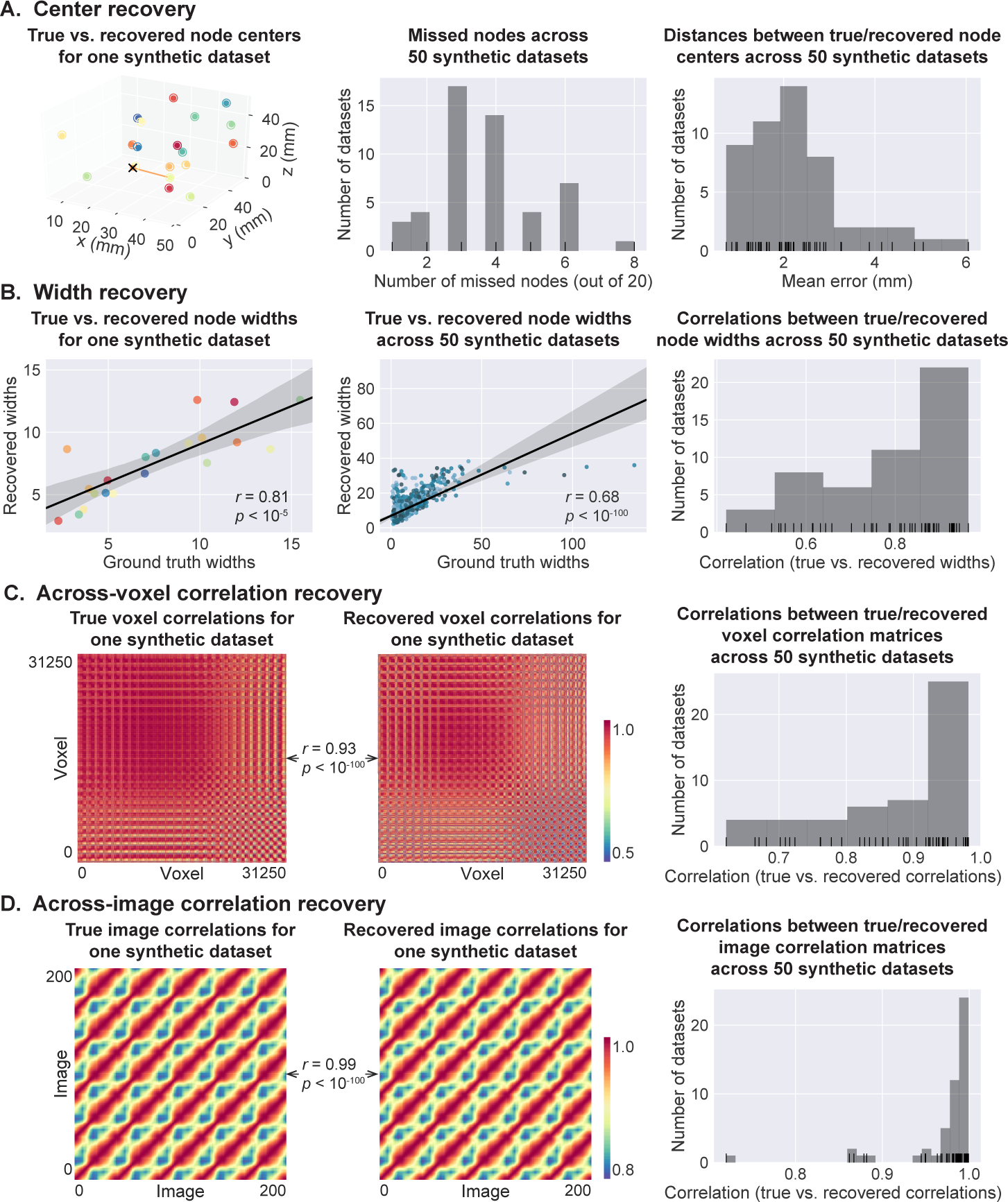
Recovered structure from 50 synthetic datasets. A. Center recovery. *Left*. Each color denotes a different node’s RBF in the global template of one synthetic dataset. The filled circles indicate the true locations of the RBF centers, and the open circles indicate the RBF centers recovered by HTFA. To facilitate visual comparison between the true and recovered locations, we have drawn a line between each recovered RBF center and the closest matching RBF center in the synthetic template. One “missed” RBF that was not assigned any nodes by HTFA (as determined by using this matching technique) is denoted by a black ×. *Middle*. Number of missed nodes across 50 synthetic datasets. *Right*. Average distances between the true and recovered (global) node centers, across 50 datasets. The averages include distances between “missed” nodes and the nearest recovered node. **B. Width recovery**. *Left*. Each dot denotes an RBF’s true and recovered (global) widths. The correlation reported in the panel is between the true and recovered RBF widths. To enable us to make comparisons in this panel, we have assigned each recovered node to the node in the original template with the closest RBF center (in terms of Euclidean distance). The colors match those in Panel A (left). *Middle*. This panel is in the same format as the left panel, but combines results across 50 synthetic datasets. The dot colors denote which dataset they came from (one color per dataset). *Right*. Correlations between true and recovered (global) widths across 50 synthetic datasets. **C. Across-voxel correlation matrices**. The left and middle panels display the true and recovered voxel-to-voxel correlation matrices for one synthetic dataset. The right panel displays a summary of the correlations between the true vs. recovered across-voxel correlation matrices across 50 synthetic datasets. **D. Across-image correlation matrices**. The left and middle panels display the true and recovered image-to-image correlation matrices for one synthetic dataset. The right panel displays a summary of the correlations between the true vs. recovered across-image correlation matrices across 50 synthetic datasets.

Our primary goal in examining the synthetic dataset was to assess whether the patterns (i.e. the RBF nodes and node weights) revealed by applying HTFA (with *K* = 20) to the dataset corresponded to the known patterns in the data. We first compared the global RBF centers and widths recovered by the model to the RBF centers and widths in the original template image. We found that the nodes’ RBF centers recovered by applying HTFA to the synthetic data closely matched the template’s RBF centers (Fig. 3A), and that the recovered RBF widths were correlated with the template RBF widths (Fig. 3B).

We next asked whether HTFA accurately recovered the correlational patterns in the synthetic data. We found that both the mean voxel-to-voxel correlation matrices (Fig. 3C) and the mean across-image correlation matrix (Fig. 3D) recovered by HTFA were strongly correlated with the true patterns we embedded into the synthetic data. (Recovery statistics for one example dataset, along with summaries of how well each pattern was recovered across all 50 datasets, are reported in the figure.) Taken together, these analyses show that HTFA is able to accurately infer (from synthetic data) the locations and sizes of the underlying RBF nodes, the correlation patterns within the images, and the correlations across images. We next turn to a series of analogous analyses on fMRI data.

### Case study 2: full-brain network dynamics are modulated by story listening

We examined fMRI data collected as participants listened to an audio recording of a story (*intact* condition; 36 participants), listened to time-scrambled recordings of the same story (18 participants in the *paragraph-scrambled* condition listened to the paragraphs in a randomized order and 25 in the *word-scrambled* condition listened to the words in a randomized order), or lay resting with their eyes open in the scanner (*rest* condition; 36 participants). We sought to demonstrate how HTFA may be used to efficiently discover and examine dynamic functional connectivity patterns in (real) multi-subject fMRI datasets. This story listening dataset was collected as part of a separate study, where the full imaging parameters, image preprocessing methods, and experimental details may be found [32]. The dataset is available <u>here</u>.

In contrast to the synthetic data we examined in Case Study 1, in real datasets there are no “ground truth” parameter values to compare to the recovered estimates. Instead, we sought to explore how well the patterns HTFA discovered could be used to decode which specific moments in the story participants were listening to. We also sought to explore whether (and how) decoding performance varied with the properties of the stimuli the participants experienced (see *Materials and methods* for a description of the decoding procedure). We first used a cross-validation procedure to determine the optimal number of nodes for efficiently representing the data while still capturing the relevant structure (see *Materials and methods*). We used all of the data from all of the experimental conditions (intact, paragraph-scrambled, word-scrambled, and rest) in this procedure-in other words, all of the experimental conditions were effectively lumped together into a single dataset. The analysis indicated that *K* = 700 nodes was optimal.

The fitted model provided estimates for each participants’ node locations, widths, and time-varying activities. Further, because the global model connects the subject-specific models, we were able to compare different participants’ activity patterns, even if their underlying nodes were not in exactly the same locations. We reasoned that, on one hand, this aspect of HTFA might improve our ability to capture cognitively relevant patterns (relative to examining the voxel activities directly). On the other hand, representing the brain images in the lower-dimensional space captured by HTFA necessarily results in information loss relative to the original voxel activity data. Effectively, HTFA blurs out high spatial frequency details from the images, where the precise amount of blurring depends on how large the nodes are and how many nodes there are overall.

We next evaluated how well HTFA is able to capture cognitively relevant brain patterns. We performed a decoding analysis, using cross-validation to estimate (using other participants’ data) which parts of the story each HTFA-derived brain activity pattern corresponded to (see *Materials and methods*). We note that our primary goal was not to achieve perfect decoding accuracy, but rather to use decoding accuracy as a benchmark for assessing whether different neural features specifically capture cognitively relevant brain patterns.

Separately for each experimental condition, we divided participants into two groups. We then computed the average activity for each group, for each of 241 overlapping 90 s (60 TR) time windows. (The 90 s window length we used in our analyses followed [32].) This yielded one activity pattern for each group of participants, for each time window. Next, for each time window, we correlated the group 1 activity patterns in that window with the group 2 activity patterns. Using these correlations, we labeled the group 1 timepoints using the group 2 timepoints with which they were most highly correlated; we then computed the proportion of correctly labeled group 1 time windows. (We also performed the symmetric analysis whereby we labeled the group 2 timepoints using the group 1 timepoints as a template.) We repeated this procedure 100 times (randomly re-assigning participants to the two groups each time) to obtain a distribution of decoding accuracies for each experimental condition. (There were 241 time windows, so chance performance on this decoding test is 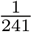.)

As a baseline, we first used the participants’ full-brain voxel activity patterns (44,415 voxels; **Y**_1…*S*_) to decode story timepoints. These voxel-based decoders achieved reliably above-chance performance on data from all four experimental conditions (*ts*(99) > 22, *ps*< 10^−400^; Fig. 4), with the best average performance in the intact condition (11.2% accuracy) and the worst average performance in the rest condition (1.5% accuracy). This shows that the original data (that we applied HTFA to) contained information about the story times that our correlation-based decoders could pick up on. Our finding that the decoders performed slightly (but reliably) above-chance during rest was unexpected. This may reflect reliable changes either in participants’ attentional states or properties of the BOLD signal over the course of the resting state scans.

We compared the performance of the voxel-based decoders to the decoding performance achieved using decoders trained on the HTFA node weights - i.e. the inferred timepoint-by-timepoint activities of the 700 nodes derived for each participant (**W**_1…*S*_). Like the voxel-based decoders, these node activity-based decoders achieved reliably above-chance performance on data from all four experimental conditions (*ts*(99) > 21, *ps*< 10^−40^; Fig. 4A). The decoding performance of the node-based and voxel-based decoders were similar (left two bars of each group in Fig. 4A; decoding accuracies match within 1%) and neither performed reliably better. We also ran analogous decoding analyses using parcel activities derived from resting state fMRI data [31] and factor weights derived from ICA with dual regression [2]. Each of these activity-based decoders performed similarly (maximum difference in decoding accuracy across all experimental conditions and methods: 2.2%); for detailed comparison statistics see Table S1.

Although the above analysis shows that HTFA node activities achieve decoding performance similar to voxel activities, the real strength of HTFA is in its enabling efficient computations that involve dynamic functional connectivity patterns (whereas these computations are in some cases intractable in the original voxel space). Following the logic of [32], we reasoned that brain activities during story listening should capture two sources of information. First, some activity should reflect the story itself. Because every participant (within each condition) listened to the same stimulus, this story-driven activity should be similar across people. Second, some activity might reflect idiosyncratic thoughts or physiological processes specific to each individual, independent of the story. This non-story-driven activity should not be similar across people. To home in on the former (story-driven) contribution to functional connectivity, we used ISFC [32]. This approach measures the correlations between brain regions of *different individuals* in each of several sliding windows (see *Materials and methods* for details). The resulting ISFC patterns are analogous to standard within-brain functional connectivity patterns (which reflect the correlational structure, across brain regions, within an individual’s brain), but they should reflect stimulus-driven activity. Decoders trained and tested on these HTFA-derived ISFC patterns achieved reliably above-chance performance on data from all experimental conditions (*ts*(99) > 10, *ps*< 10^−15^; Fig. 4A). Although these ISFC-based decoders were out-performed by the voxel-based and node activity-based decoders, it is important to note that the correlation-based features that the ISFC-based decoders utilize are fundamentally different than node activity patterns. We carried out analogous analyses using the parcel activity-based decoders and ICA-based factor activation-based decoders described above. The parcel-based analysis yielded similar results to what we observed using HTFA (whereby ISFC-based decoders performed reliably worse than activity-based decoders), and the ICA-based analyses yielded no significant differences between ISFC-based and activity-based decoders (Tab. S1).

Given the above results, we wondered whether the node activity-based decoders and ISFC-based decoders might be picking up on partially non-overlapping sources of cognitively relevant information. If so, decoders trained on a mix of activity-based and ISFC-based features might outperform decoders trained on only a single class of features.

We therefore designed a fourth set of decoders whose predictions were a 50-50 mix of the node activity-based decoders and the ISFC-based decoders; see *Materials and methods* for additional details. These hybrid decoders reliably out-performed the other decoders we examined on all experimental conditions except the rest condition (intact, paragraph, and word conditions: *ts*(99) > 14, *ps*< 10^−8^; Fig. 4A). This finding is consistent with the notion that activity-based and ISFC-based patterns contain *different* information about the story moments people were listening to. We replicated this finding that a blend of activity-based and ISFC-based decoders outperform solely activity-based or ISFC-based decoders using atlas-derived parcel activations and ICA factor weights (Fig. 4A, Tab. S1).

To follow up on this result, we set out to determine the optimal mixing proportion of the two types of HTFA-derived features. We defined a mixing parameter, *ϕ*, where *ϕ* = 0 corresponds to activity-based decoding; *ϕ* = 1 corresponds to ISFC-based decoding; and 0 < *ϕ* < 1 reflects a weighted mixture of activity-and ISFC-based decoding. We then re-ran the “mixture” decoding analysis in Figure 4A using 21 linearly spaced values of *ϕ* ranging between 0 and 1, inclusive. As shown in Figure 5A, the decoding accuracy (for the intact condition) peaked for *ϕ* = 0.25, corresponding to a mix of 75% activity-based and 25% ISFC-based features.

These analyses indicate that HTFA enables the efficient implementation of more sophisticated decoders that would otherwise be computationally expensive. These decoders (that incorporate correlational information, or a mix of activity-based and correlation-based information) out-perform decoders trained only on raw voxel activity patterns. Further, HTFA yields similar decoding performance to other established methods (see Tab. S1 for detailed comparisons).

### Case Study 3: full-brain networks are modulated by movie viewing

As a secondary test of our results from Case Study 2, we ran an analogous set of analyses on data collected as 17 participants viewed an episode from the BBC television show *Sherlock* [9]. (Experimental and imaging methods may be found in [9].) As in Case Study 2, we used a cross validation procedure to determine the optimal number of network nodes for the dataset (see *Materials and methods*). This procedure revealed that K = 900 nodes were optimal. We then applied HTFA to the full dataset, which comprised (for each participant) 43,371 voxel activities over 1976 TRs. The dataset is available <u>here</u>.

As shown in Figure 4B, all 10 classes of features we examined in Case Study 2 could also be used for this movie viewing dataset to reliably decode which timepoint participants were viewing (Fig. 4B; *ts*(99) > 78, *ps*< 10^−90^). Following our approach in Case Study 2, we divided the experiment into 60 TR sliding windows. Therefore chance performance on this test is 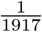 (1976 - 60 + 1 = 1917 unique sliding windows).

We next carried out a series of analyses using HTFA, atlas-based parcels, and ICA factors to determine which aspects of the activity and connectivity patterns yielded the best decoding accuracy. As with Case Study 2, we found that mixed activity-ISFC decoders outperformed solely activity-based or ISFC-based decoders. (For detailed performance statistics see Table S2.)

We also carried out an exploratory analysis to estimate the optimal mixing proportions of HTFA node activity-based and ISFC-based features (Fig. 5B). The decoders with *ϕ* = 0.3 (reflecting a mixture of 70% activity-based and 30% ISFC-based features) performed best. These analyses provide additional evidence that the activity and connectivity patterns contain partially non-overlapping information about the moments in the movie the participants were viewing.

## Discussion

We proposed HTFA, a probabilistic approach to discovering and examining full-brain patterns of dynamic functional connectivity in multi-subject fMRI datasets. In Case Study 1, we used a synthetic dataset to demonstrate HTFA’s ability to recover known patterns in synthetic data, and in Case Studies 2 and 3 we applied HTFA to real data, and showed that the resulting patterns could be used to decode story listening and movie viewing times. We also compared the performance of HTFA-derived decoders to decoders trained on features from other established methods, and we obtained similar decoding results.

### Benefits and costs of our approach

HTFA was designed to provide an efficient means of examining dynamic full-brain connectivity patterns in multi-subject datasets. What would it have taken to study the same ISFC patterns we examined in Case Study 2 using voxel-based methods? After masking out non-gray matter voxels and warping every subject’s data to a common brain space (see *Materials and methods*), each fMRI volume comprises 44,415 voxels. Therefore each voxelwise ISFC matrix contains roughly 986 million unique entries (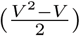). Assuming single precision storage (4 bytes per number), the ISFC matrix for each timepoint would require roughly 3.9GB of memory. By contrast, each HTFA-derived ISFC matrix for 700 nodes requires fewer than 250 thousand unique entries (1 MB). In general, for decoders that combine activity-based and ISFC-based features, the computational savings (in terms of representational compactness) gained by using HTFA-derived networks (of *K* nodes) rather than the original voxel data is given by 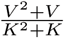. For our Case Study 2, the HTFA-derived ISFC analyses in Figure 4A were approximately 4,000 times more computationally efficient than the corresponding voxel-based analyses would have been. Following this logic, we achieved a similar computational efficiency savings for Case Study 3; Fig. 4B. We note that these representational compactness benefits also apply to other dimensionality reduction approaches (e.g. see *Introduction*). Although HTFA and other dimensionality reduction approaches facilitate efficient computations of full-brain correlation patterns, we note that with sufficient computing power, optimized approaches have been developed for computing these matrices directly [37].

Another benefit of HTFA’s separation from voxel space is its ability to naturally fill in missing observations (a property we exploit to determine the optimal number of nodes). Techniques like probabilistic PCA [34] can fill in missing voxel activities using the data covariance matrix, provided that we observe at least *some* activities from those missing voxels (in other images). However, suppose that *all* activities from a given voxel were missing-or more realistically, suppose that we wish to estimate what the activities would have been at any arbitrary point in space. Because PCA does not explicitly represent the voxels’ spatial locations, neither PCA nor probabilistic PCA can accurately predict activity patterns at these never-observed voxels. HTFA, by contrast, naturally predicts the missing data by simply evaluating each node’s RBF at the corresponding location in space.

These missing data examples also provide insights into other benefits of allowing nodes to exist in real space rather than considering only the set of voxel locations. For example, HTFA allows for different subjects’ data to be sampled at different resolutions, or to contain different numbers of voxels. In principle, different subjects’ data may even come from different recording modalities (e.g. one subject may contribute fMRI data and another may contribute EEG data). In this way, HTFA provides a common framework for describing neural data in general that transcends the specifics of the recording (modality, spatial or temporal resolution, etc.). For additional discussion of the benefits of spatial-based (rather than voxel-based) nodes see [18], and for an example of how similar models may be used to analyze EEG data see [19].

Note that using HTFA to examine connectivity patterns may not always out-perform voxel-based approaches. In particular, to the extent that the relevant patterns are high spatial frequency (at the level of single voxels), those patterns will be better described by voxel-based approaches than RBF nodes. (Representing brain images as sums of RBFs effectively blurs out the images in space, where the amount of blurring is inversely proportional to the number of nodes.) To address this issue, we developed an algorithm for estimating the minimum number of nodes required to reliably describe the data up to a desired level of precision (see *Materials and methods*).

Although we have focused in this manuscript on highlighting the strengths of HTFA in particular, it is worth noting the strengths of related approaches like the atlas-based segmentation and ICA with dual regression approaches that we compared with HTFA in Case Studies 2 and 3. For example, atlas-based segmentation methods obey anatomical boundaries well. This property is especially useful in applications aimed at discovering or exploring specific anatomical features or patterns. ICA-based approaches yield statistically independent features. This property is especially useful in applications that leverage or assume statistical independence between difference neural sources. HTFA, in turn, yields spatially compact features (which are useful for network visualizations and explorations) that partially overlap. This latter property may help detangle the contributions of spatially overlapping, but functionally distinct, activity patterns.

### Concluding remarks

HTFA provides an efficient means of examining dynamic full-brain functional connectivity patterns, thereby making it easier to study how connectivity patterns relate to the cognitive processes they support. Further, HTFA’s compact representations of connectivity facilitate studying connectivity with algorithms that are too expensive to use on the original data.

## Acknowledgements

We acknowledge useful discussions with Jonathan Cohen, Janice Chen, Justin Hulbert, J. Benjamin Hutchinson, Talia Manning, Peter Ramadge, Erez Simony, and Nicholas Turk-Browne. This work was supported by the NSF/NIH Collaborative Research in Computational Neuroscience Program, grant number NSF IIS-1009542; NSF EPSCoR Award Number 1632738; and a grant from the Intel Corporation. The content is solely the responsibility of the authors and does not necessarily represent the official views of our supporting organizations. The implementation of HTFA that we used in this paper may be found in the <u>BrainIAK Toolbox</u> [8].

## supplemental materials for A probabilistic approach to discovering dynamic full-brain functional connectivity patterns

### Overview

These supplemental materials provide additional details about HTFA and the decoding results reported in Case Studies 2 and 3 in the main text.

### Formal definition and notation

We formulate HTFA as a probabilistic latent variable model, which can be represented in graphical model notation. In the graphical model (Fig. 2, main text), variables associated with the subject-specific template are found in the yellow plate. These include the subject-specific RBF centers (*μ*_1…*K*,1…*S*_), RBF widths (*λ*_1…*K*,1…*S*_), and per-image factor weights (*w*_1…*N*,1…*K*,1…*S*_), as well as the observed images (***y***_1…*N*,1…*S*_). Variables associated with the global template are found outside of the yellow plate; these include the global RBF centers 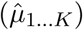 and the global RBF widths 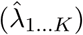. The subject-specific templates are conditioned on the global template, thereby associating data from different subjects. (This interaction between the subject-specific and global templates occurs where the yellow and blue plates overlap.)

**Figure 2.**
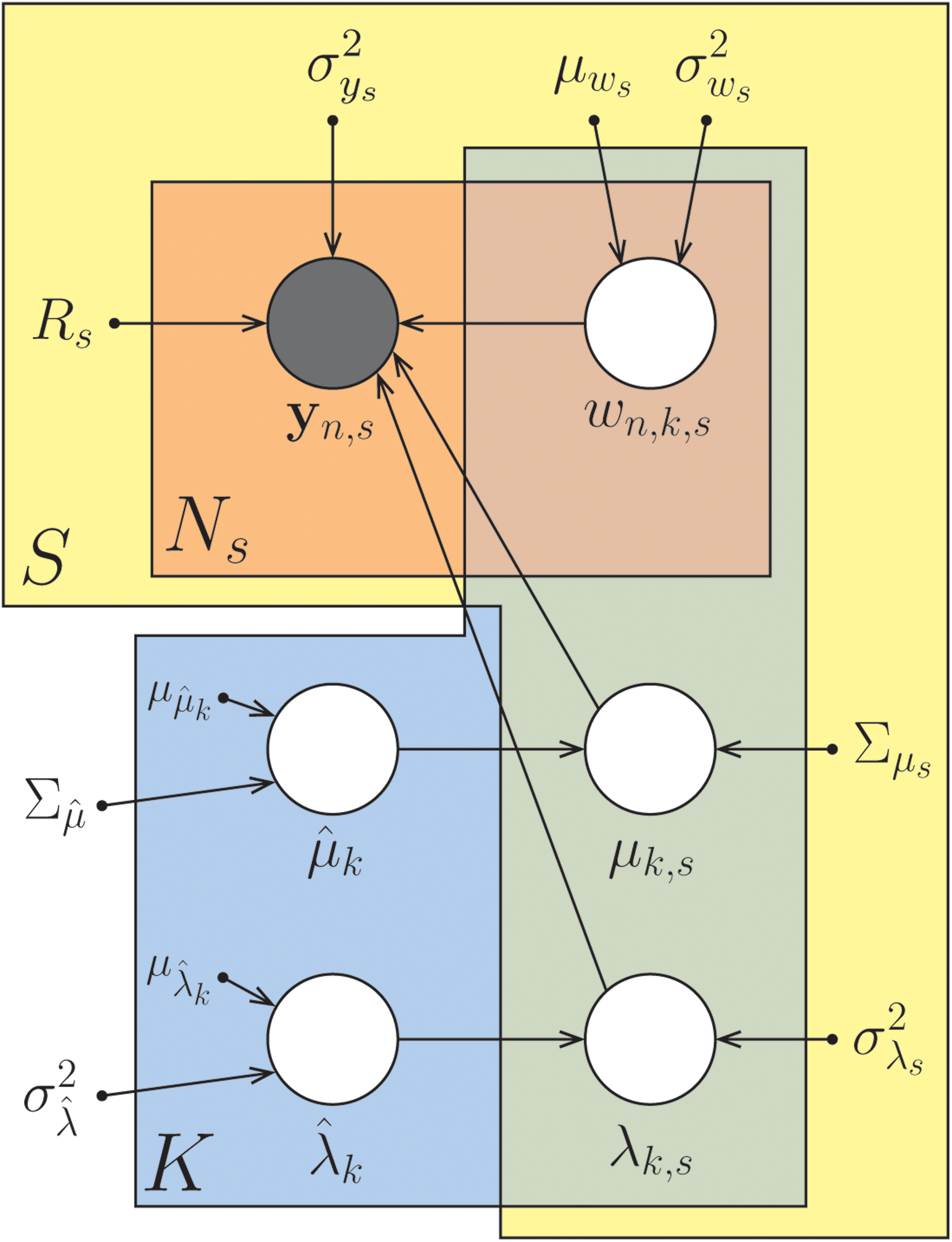
Graphical model for HTFA. Each variable in the model appears as a circle; hidden variables are unshaded and observed variables are shaded. The variables include **y**_*n,s*_ (observed image *n* from subject *s*); *w*_*n,k,s*_ (node *k*’s weight, or activity, in image *n* from subject *s*); *μ*_*k,s*_ (node *k*’s center coordinate for subject *s*); *λ*_*k,s*_ (node *k*’s width for subject *s*); 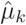*k* (node *k*’s center in the global template); and 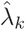_*k*_ (node *k*’s width in the global template). Hyperparameters (defined in the *supplemental materials*) are denoted by dots. Arrows denote conditional dependence, originating at terms that appear on the right sides of conditionals and pointing towards terms that appear on the left sides. Rectangular plates denote repeated structure, where the number of copies is indicated within each plate: *N*_*s*_ (number of images from subject *s*); *S* (number of subjects), and *K* (number of nodes). For a comprehensive introduction to graphical models see [4].

The structure of the graphical model specifies the conditional dependencies in HTFA:

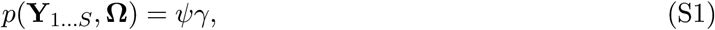

where **Y**_*s*_ is the *N*_*s*_ by *V*_*s*_ matrix of images from subject *s*,^1^ **Ω** is the set of hidden variables in the model

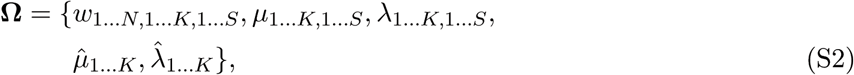

the probability of the set of subject-specific templates *ψ* is

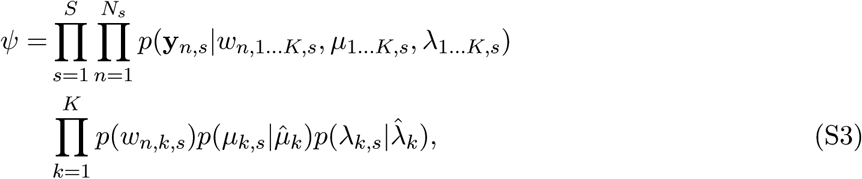

and where the probability of the global template *γ* is

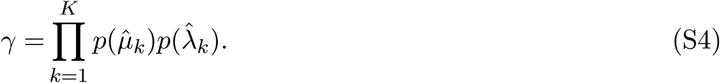

(Note that the hyperparameters have been omitted to simplify the notation; see Algorithm S1 for the full specification.)

Another useful way to describe HTFA is through the *generative process* it implies. HTFA’s generative process is an algorithm that, when run, generates a single sample from HTFA’s joint distribution (Eqn. S1), yielding one value for each hidden variable and a multi-subject fMRI dataset. HTFA’s generative process is detailed in Algorithm S1. The generative process starts by drawing a set of global RBF parameters, and from there draws subject-specific RBF parameters and factor weights, and from there draws the data. We emphasize that this “algorithm” represents the imaginary generative process from which the model assumes the data arises. When we fit the model by applying HTFA to a dataset, we “reverse” the generative process by starting with each subject’s data, which we use to estimate their subject-specific RBFs and weights, which we use in turn to estimate the global RBFs.

#### Algorithm S1: HTFA’s generative process

Here **F**_*s*_ is the *K* by *V*_*s*_ factor image matrix for subject *s*, which depends on their subject-specific RBF centers (*μ*1…*K,s*), RBF widths (*λ*1…*K,s*), and their *V*_*s*_ by 3 voxel location matrix (**R**_*s*_).

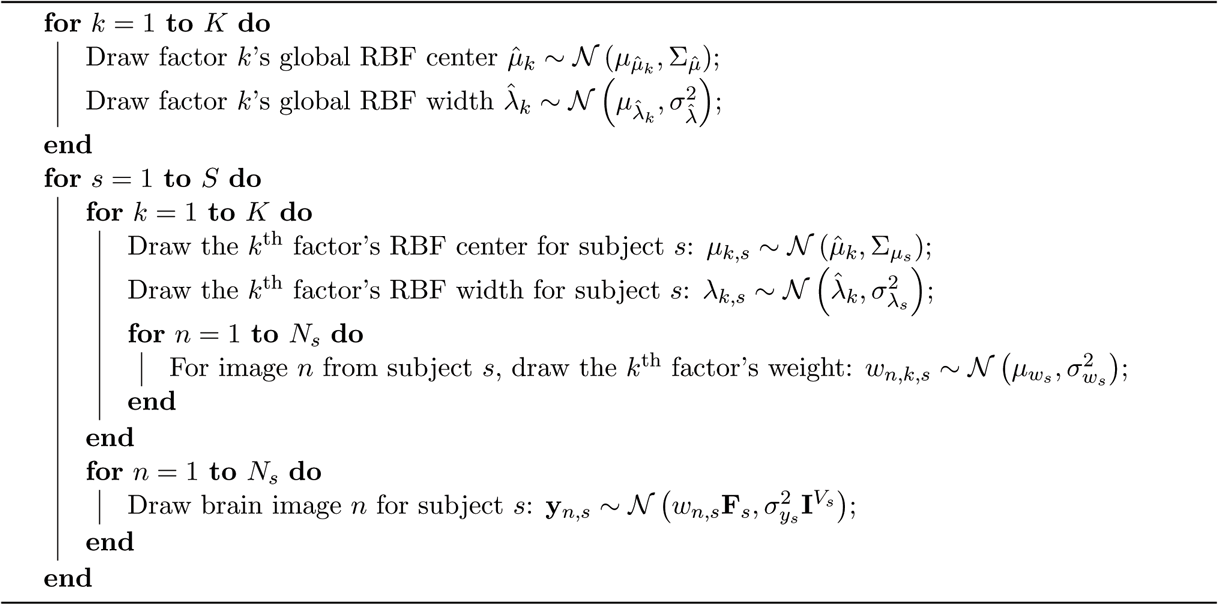

Our goal in applying HTFA to a dataset is to compute the probability distribution over the model’s hidden variables (e.g. RBF centers and widths, and factor weights) given the dataset. The posterior distribution *p*(**Ω**|**Y**_1…*S*_) tells us how likely each hidden variable is to be set to a particular value, given the data and our prior assumptions about what these values should be. In theory, we could use Bayes’ rule to compute this posterior:

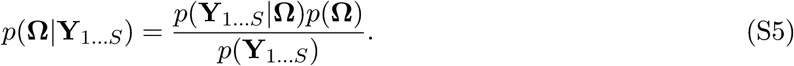

However, computing the denominator (as for most models) is intractable:

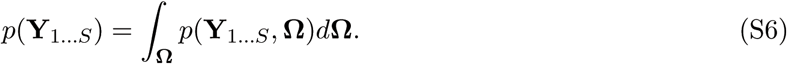

Notice that solving for *p*(**Y**_1…*S*_) requires integrating over all possible values of each of the hidden variables in the model, for which there is no analytic solution. Therefore, instead of computing the full posterior distribution, here we have developed an efficient *maximum a posteriori* (MAP) algorithm for estimating the most probable values of the hidden variables under the posterior. We provide a high-level description of how we can use MAP inference to efficiently apply HTFA to large multi-subject fMRI datasets in *Materials and methods*; a full description may be found in [1]. We have also released an open source toolbox for applying HTFA to fMRI data, available for download <u>here</u>.

### Detailed decoding results

Tables S1 and S2 provide detailed comparisons between the decoding performance obtained using decoders trained using all of the different features (voxel activity, HTFA node activity, atlas-parcel activity, and ICA with dual regression factor weights) and post-processing steps (raw activity, ISFC, and a 50-50 mix of activity and ISFC). The details of the decoding analyses may be found in the main text (*Materials and methods*).

**Table SI.**
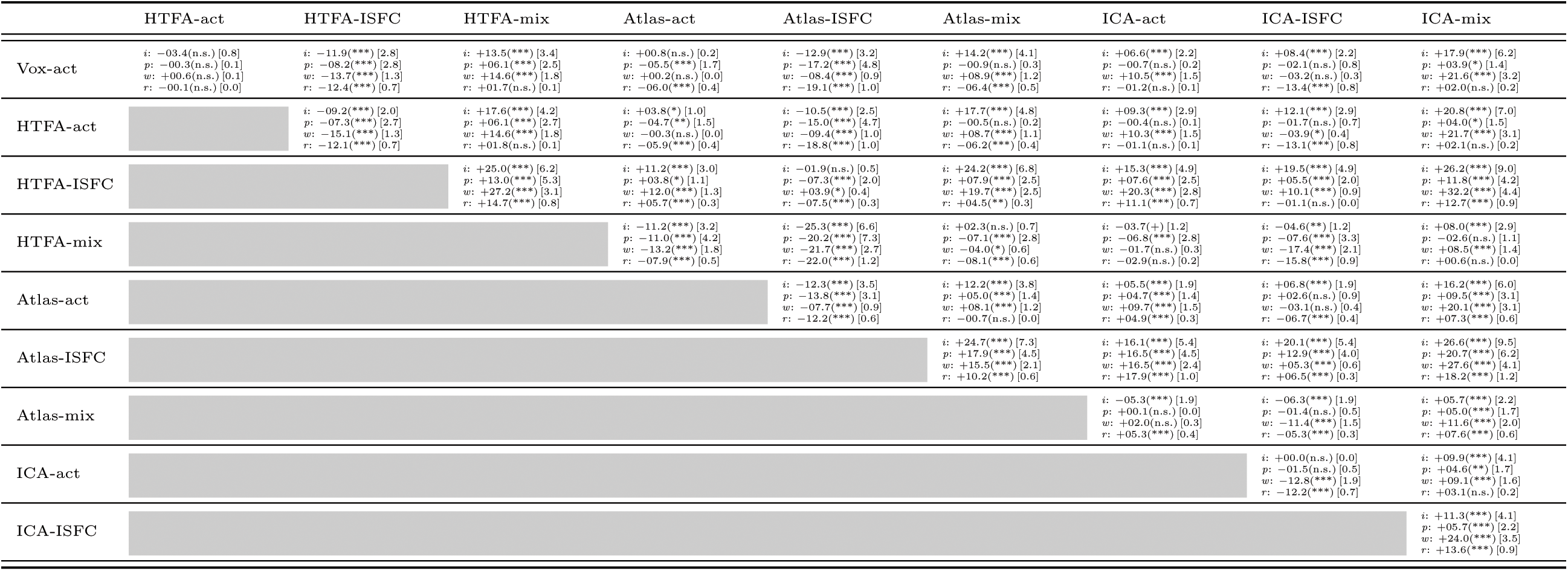
Each cell displays the results of *t*-tests comparing the (bootstrap-estimated) decoding accuracy observed using two decoders, for each of the four experimental conditions: *i*ntact, *p*aragraph-scrambled, *w*ord-scrambled, and *r*est. All *t*-tests have 99 degrees of freedom (number of bootstrap runs minus 1); the first numbers on each line are the *t*-values, and the corresponding Bonferroni-corrected significance values are in parenthesis (* * *: *p* < 0.001; **: *p* < 0.01; *: *p* < 0.05; +: *p* < 0.1; n.s.: *p* ≥ 0.1). The numbers in square brackets report the mean absolute difference in percent decoding accuracy. Positive *t*-values reflect decoders trained on the column’s feature outperforming the decoders trained on the row’s feature. A total of 10 types of features are compared: voxel activity (Vox-act), HTFA node activity (HTFA-act), HTFA node ISFC (HTFA-ISFC), HTFA node activity-ISFC mixture (HTFA-mix), Atlas-based parcel activity (Atlas-act), Atlas-based parcel ISFC (Atlas-ISFC), Atlas-based parcel activity-ISFC mixture (Atlas-mix), ICA factor activity (ICA-act), ICA factor ISFC (ISFC-ISFC), and ICA factor activity-ISFC mixture (ICA-mix).

**Table S2.** Each cell displays the results of *t*-tests comparing the (bootstrap-estimated) decoding accuracy observed using two decoders. All *t*-tests have 99 degrees of freedom (number of bootstrap runs minus 1); the first numbers on each cell are the *t*-values, and the corresponding Bonferroni-corrected significance values are in parenthesis (* * *: *p* < 0.001; **: *p* < 0.01; *: *p* < 0.05; +: *p* < 0.1; n.s.: *p* ≥ 0.1). The numbers in square brackets report the mean absolute difference in percent decoding accuracy. Positive *t*-values reflect decoders trained on the column’s feature outperforming the decoders trained on the row’s feature. A total of 10 types of features are compared: voxel activity (Vox-act), HTFA node activity (HTFA-act), HTFA node ISFC (HTFA-ISFC), HTFA node activity-ISFC mixture (HTFA-mix), Atlas-based parcel activity (Atlas-act), Atlas-based parcel ISFC (Atlas-ISFC), Atlas-based parcel activity-ISFC mixture (Atlas-mix), ICA factor activity (ICA-act), ICA factor ISFC (ISFC-ISFC), and ICA factor activity-ISFC mixture (ICA-mix).

|  | HTFA-act | HTFA-ISFC | HTFA-mix | Atlas-act | Atlas-ISFC | Atlas-mix | ICA-act | ICA-ISFC | ICA-mix |
| --- | --- | --- | --- | --- | --- | --- | --- | --- | --- |
| Vox-act | -13.7(***) [1.6] | -09.6(***) [1.1] | +44.7(***) [5.2] | +02.5(n.s.) [0.3] | -22.9(***) [2.7] | +41.1(***) [5.8] | -16.5(***) [2.3] | +27.1(***) [3.2] | +36.3(***) [5.3] |
| HTFA-act |  | +04.2(**) [0.5] | +60.0(***) [6.8] | +14.7(***) [1.9] | -09.6(***) [1.1] | +53.3(***) [7.4] | -05.3(***) [0.7] | +41.6(***) [4.8] | +48.1(***) [6.8] |
| HTFA-ISFC |  |  | +55.6(***) [6.4] | +11.1(***) [1.4] | -13.7(***) [1.5] | +49.8(***) [6.9] | -08.7(***) [1.2] | +37.4(***) [4.3] | +44.7(***) [6.4] |
| HTFA-mix |  |  |  | -36.9(***) [4.9] | -68.7(***) [7.9] | +04.2(**) [0.6] | -54.6(***) [7.5] | -17.1(***) [2.0] | +00.1(n.s.) [0.0] |
| Atlas-act |  |  |  |  | -22.8(***) [3.0] | +35.4(***) [5.5] | -17.3(***) [2.6] | +21.5(***) [2.9] | +31.2(***) [4.9] |
| Atlas-ISFC |  |  |  |  |  | +60.6(***) [8.5] | +02.6(n.s.) [0.4] | +50.4(***) [5.9] | +55.3(***) [7.9] |
| Atlas-mix |  |  |  |  |  |  | -50.9(***) [8.1] | -18.3(***) [2.6] | -03.4(*) [0.6] |
| ICA-act |  |  |  |  |  |  |  | +39.6(***) [5.5] | +46.5(***) [7.6] |
| ICA-ISFC |  |  |  |  |  |  |  |  | +14.0(***) [2.0] |

## Footnotes

1 Note that both the number of images from subject s, *N*_*s*_, and the number of voxels from subject s, *V*_*s*_, may vary across subjects.

